# Sex-Specific DNA-Replication In The Early Mammalian Embryo

**DOI:** 10.1101/2023.12.22.572993

**Authors:** Jason Alexander Halliwell, Javier Martin-Gonzalez, Adnan Hashim, John Arne Dahl, Eva Ran Hoffmann, Mads Lerdrup

## Abstract

The timing of mammalian DNA replication is crucial for minimizing errors and is influenced locally by genome usage and chromatin states. However, our understanding of replication timing in the unique environment that exists in newly formed mammalian embryos is limited. Here, we performed genome-wide investigations of replication timing in mouse zygotes and 2-cell embryos. We discovered that zygotes lack a conventional replication timing, but a program emerged in 2-cell embryos. Notably, this program differs from embryonic stem cells and shows asynchrony between the parental genomes. We observed late replication of maternal pericentromeric regions and DNA associated with the nuclear lamina in each parental genome, whereas unexpected early replication occurred in regions with histone marks deposited by Polycomb Repressive Complexes on the maternal chromatin. This atypical and asynchronous replication of the two parental genomes may advance our understanding of replication stress in early human embryos and trigger strategies to reduce errors and aneuploidies.

## Main

DNA replication is a fundamental process ensuring accurate transfer of genetic information when cells divide. Its precise orchestration follows a well-defined temporal order, known as the replication timing program^1,2^. Replication timing has emerged as a key cellular fingerprint that is associated with various genomic features, such as gene density, gene transcription, histone modifications, DNA methylation, and 3D genome organization^3–8^. Furthermore, it exhibits a remarkable level of synchrony across single cells and their homologous chromosomes^9,10^.

Significant advancements in our comprehension of replication timing have been facilitated by a range of sequencing technologies collectively referred to as Repli-seq^9–11^. Repli-seq examines copy number variations of S phase cells and normalizes them against G1 reference cells. More recently, the advent of single-cell sequencing approaches has revolutionized the investigation of Replication timing in individual cells^9,10,12^.

Despite significant progress, our comprehension of replication timing heavily relies on established replication programs derived from *in vitro* cell lines. However, little is known about how *de novo* replication programs are established after fertilization in the developing embryo.

Human and mouse oocytes and embryos are characterized by several transformative changes in their epigenetic landscapes^13–17^, and a permissive transcriptional environment that allows for unusually high retrotransposon activity and the emergence of alternative retrotransposon-initiated transcripts^18–21^. As the paternal genome is unpacked from the fertilizing sperm, its protamines are replaced by histones, and both the maternal and paternal genomes must exit the transcriptionally silenced states that existed in the gametes to initiate embryonic genome activation^22^, while undergoing major morphological changes^23^. Collectively, these factors could place the preimplantation genome under considerable exertion, and might contribute to the reported instances of replication stress during the first cellular divisions in mouse and human embryos^24,25^. Consequently, exceptional factors might influence and govern the replication and its timing at the early stages of mammalian embryo development.

## Results

### Single-embryo Repli-seq reveals atypical replication timing in zygotes and 2-cell embryos

To better understand the mechanisms underlying the establishment of replication timing during early mammalian embryogenesis, we developed a single embryo Repli-seq technique to study replication timing in the zygote and 2-cell stages of mouse embryos (**Fig. 1a**). The reliability of Repli-Seq relies on the precise selection of cells in S phase, therefore, we first employed EdU labeling at specific time points following fertilization. This allowed us to determine the exact timing of the zygotes entry into and exit from the first S phase (**Fig. 1a-c**). For the 2-cell embryos, we additionally used Nocodazole to arrest *in vitro* fertilized mouse zygotes in metaphase for 4 hours, aligning their cell cycles and facilitating precise selection of G1 or S phase cells based on EdU incorporation (**Fig. 1a-c**).

**Figure 1.**
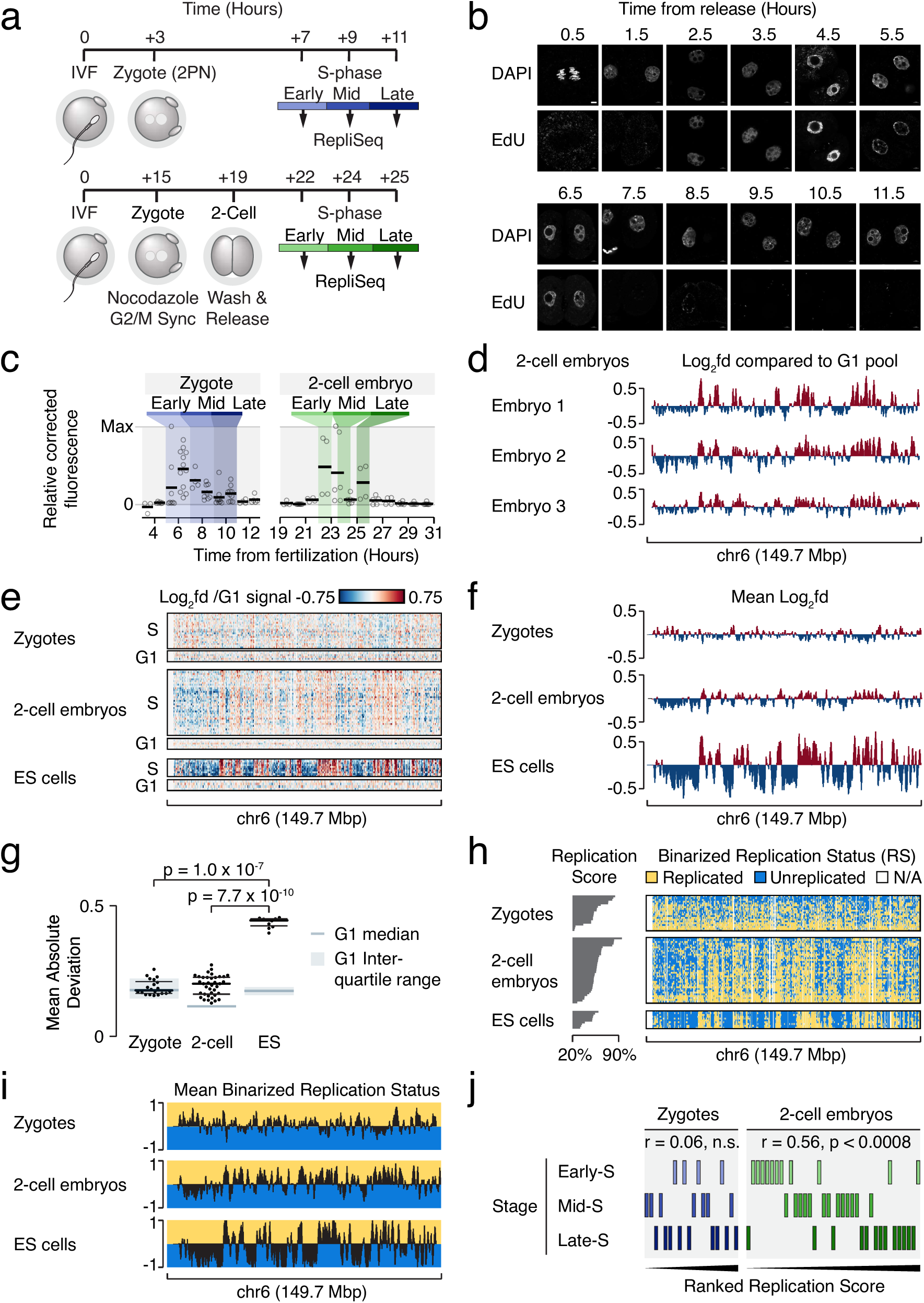
Single-embryo Repli-seq reveals atypical replication timing in zygotes and 2-cell embryos. **a)** The schematic illustrates the procedure and timing for obtaining mouse zygotes and 2-cell embryos undergoing DNA replication. In the upper panel, zygotes were selected at different times during the S phase after fertilization. In the lower panel, Nocodazole was utilized to synchronize the release of 2-cell embryos into the second cell cycle. **b)** Microscopy of EdU pulse labelling of mouse 2-cell embryos at indicated time points after release from Nocodazole-induced inhibition of mitosis. Scale bar = 5µM. **c)** Quantitation of EdU-incorporation in mouse zygotes and 2-cell embryos at indicated time points after fertilization. Coloring illustrates the time points defined as Early, Mid, and Late parts of the S-phase. **d)** Graphs showing normalized Repli-seq read densities of three S-phase 2-cell embryos at chromosome 6. Values are log2-fold differences relative to the G1-phase signal. **e, f**) The heatmap (**e**) and graph (**f**) display normalized Repli-seq read densities in S- and G1-phase mouse zygotes, 2-cell embryos, and ESCs on chromosome 6. The heatmap shows individual embryos, while the graph represents mean values. The values are presented as log2-fold differences relative to the G1-phase signal for each cell type. **g**) Beeswarm plots of the mean absolute genome-wide deviation in each Repli-seq sample. The background deviation from G1 controls for each sample is indicated with grey boxes and wide bars showing the interquartile range and median, respectively. Black bars indicate the 25, 50, and 75 percentiles of the S-phase populations. P-values obtained by two-sided Mann-Whitney U-tests Benjamini-Hochberg corrected for multiple testing. **h, i**) Heatmaps of individual binarized replication status values (**h**) and mean binarized Replication status of S-Phase mouse zygotes, 2-cell embryos and ESCs at chromosome 6 (**i**). Binarized values were derived from Repli-seq data normalized to the G1-phase signal for each cell type. The vertical order and left side bar diagram (**h**) reflect the Replication Score for each sample. **j**) Tile plot showing relationships between S-phase progression and ranked Replication Score from Repli-seq data in mouse zygotes and 2-cell embryos. r- and p-values were obtained using Spearman’s rank correlation tests Benjamini-Hochberg corrected for multiple testing.

With accurate definition of the cell cycle times, we obtained a cohort of 26 zygotes and 44 *in vitro* fertilized 2-cell stage embryos from C57BL/6N oocytes fertilized with CAST/EiJ sperm. These embryos were processed using a modified version of previously published sc-Repliseq pipelines (methods)^9,10^. To validate the reliability of our approach, we also generated replication timing profiles from individual mouse embryonic stem cells (ESCs). These replication timing profiles demonstrated a strong agreement with the previously published sc-Repli-seq profiles obtained from ESCs (**Extended Data Fig. 1a-e, Supplementary Table 1**)^9^. Based on our previous EdU determination, six, four, seven, and nine zygotes were assigned as G1, early S, mid S, and late S phase, respectively. For 2-cell embryos, seven, eleven, thirteen, and thirteen embryos were assigned to the G1, early S, mid S, and late S phase, respectively.

Our analysis of replication timing in individual 2-cell embryos revealed consistent replication timing profiles across individual samples (**Fig. 1d**). However, we observed marked changes in the overall level of differences between S phase cells and G1 cells during embryonic growth. Specifically, the zygote data exhibited a notably smaller overall deviation (**Fig. 1e,f**), while Repli-Seq profiles, generated from ESCs showed a strong and localized deviation between their S and G1 cells, which was not attributed to read depth (**Fig. 1e-g and Extended Data Fig. 1f-g, Supplementary Table 2**). 2-cell embryos displayed intermediate levels of G1/S deviation, suggesting that the changes might coincide with the embryo genome activation. While asynchronous replication between the two parental genomes is a potential cause, this could also indicate a uniform replication program that initiates from a high abundance of replication origins that are randomly distributed throughout the genome.

Notably, when we assessed the replication status of each embryo by methodology previously validated in ESCs^9^ (**Fig. 1h,i**), we found a close alignment between the assessment made by EdU incorporation (**Fig. 1b,c**) and the calculated Replication Score for each 2-cell embryos, but not for zygotes (**Fig. 1j, Supplementary Table 3**). Given the limited and insignificant differences between G1 and S phase zygotes, (**Fig. 1f,g and Extended Data Fig. 1f**), we decided to focus on the significant deviations seen in the 2-cell stage in our subsequent analyses of genome-wide localization of replication timing.

### Consistent differences in local replication timing between 2-cell embryos and ESCs

Although replication timing between our 2-cell embryos and ESCs ordered by the fraction of the genome scored as replicated ^9^ showed many similarities throughout the genome, we also observed notable local differences between the two cell types (**Fig. 2a**). These distinct differences were consistent across individual samples of the two cell types, with some displaying markedly earlier or later replication in the 2-cell embryos compared to ESCs (**Fig. 2a**).

**Figure 2.**
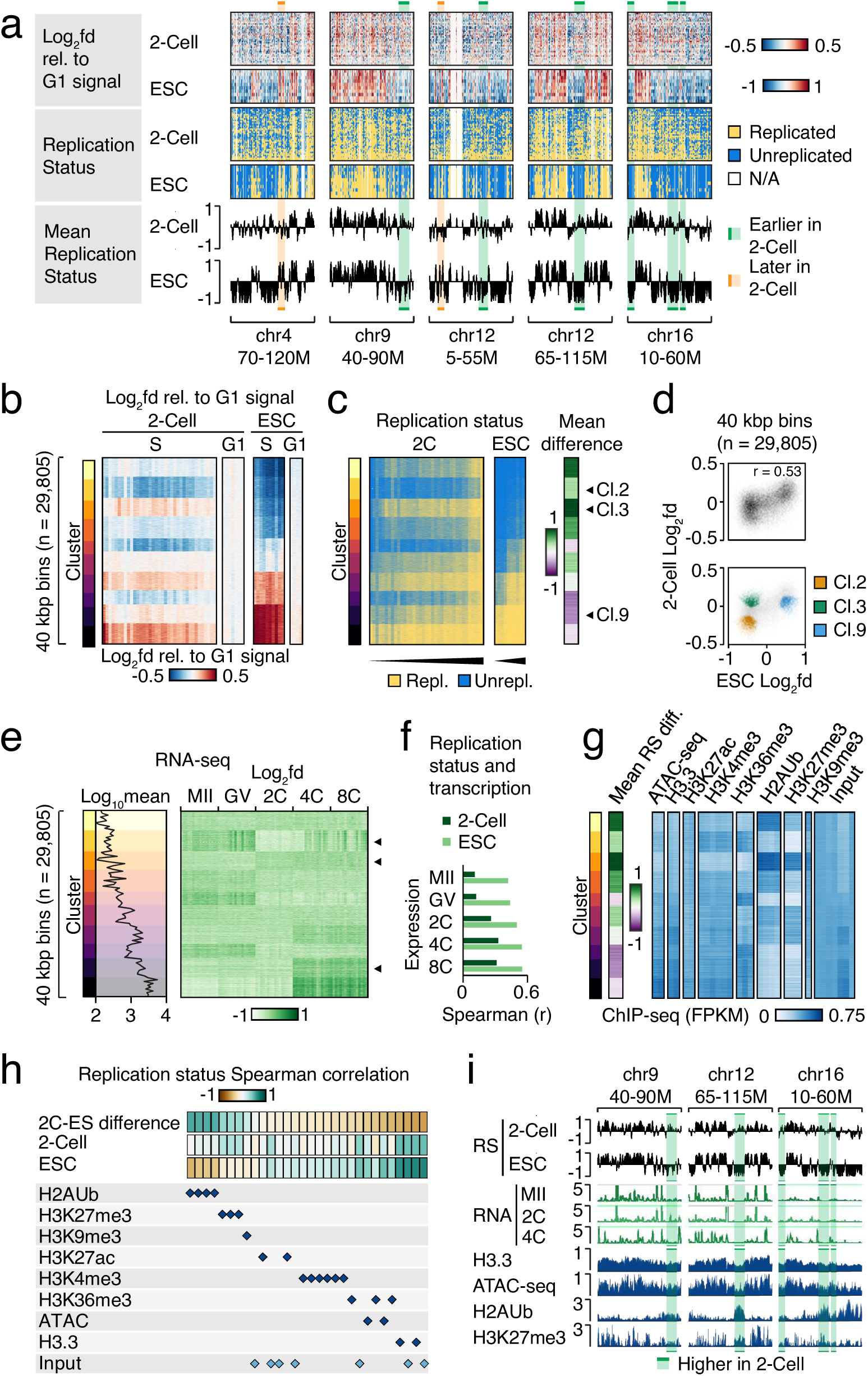
Consistent differences in local replication timing between 2-cell embryos and ESCs. **a)** Heatmaps showing normalized Repli-seq read densities (upper) and binarized Replication Status (middle) as well as graphs showing mean binarized Replication Status (bottom) of S-phase 2-cell mouse embryos and ESCs at five selected genomic loci. Orange and green highlighted regions indicate loci with marked differences between these cell types. **b)** Heatmaps showing the genome-wide signal enrichment and variation in individual Repli-seq samples from S- and G1-phase 2-cell embryos and ESCs. 40 kbp bins that contained signal from both conditions (n = 29,805) were k-means clustered according to the indicated S-phase signal from all individual 2-cell mouse embryos and ESCs. Clusters were ordered according to S-phase ESC signal. **c)** Heatmaps showing the genome-wide binarized Replication Status in individual Repli-seq samples from S-phase 2-cell embryos and ESCs ordered as in (**b**). Rightmost heatmap shows differences in the mean binarized Replication Status between 2-cell mouse embryos and ESCs. **d)** Scatter plots showing the relationship between mean normalized S-phase Repli-seq read densities in mouse ESCs (X-axis) and 2-cell embryos (Y-axis) for all 40 kbp bins (top) or three selected clusters (bottom). R-value indicates the Pearson correlation coefficient. **d)** Composite panel showing mean genome-wide transcript density (graph, left) or log2-fold differences relative to the mean (heatmap, right) from single mouse oocyte or embryo RNA-seq^19^. Signal was quantified in 40 kbp bins ordered as in (**b**). **e)** Bar diagrams showing the correlation between mean Replication Status from S-phase mouse 2-cell embryos or ES-cells and transcript density from the indicated oocyte and embryo stages. The signal was analyzed in 40 kbp bins and correlation was measured using Spearman rank correlation tests. **f)** Heatmaps showing genome-wide histone marks, histone variant, and chromatin accessibility signal from mouse oocytes. Signal was quantified in 40kbp bins ordered as in (**b**) and FPKM-normalized. Leftmost heatmap shows mean Replication status differences as in (**c**). **g)** Tile plot showing the genome-wide correlation of histone marks, histone variant, and chromatin accessibility data in mouse oocytes (indicated by lowermost blue rhombs) relative to mean Replication Status from S-phase mouse 2-cell embryos, ES-cells, and the difference between these two. The signal was analyzed in 40 kbp bins and correlation measured using Spearman rank correlation tests. **h)** Genome-browser tracks of mean binarized Replication Status (top, black), RNA-seq (middle, green), histone mark and chromatin accessibility (bottom, blue) at example loci with relatively early 2-cell replication (highlighted in green).

To assess these differences in an unbiased manner and on a global scale, we performed k-means clustering of the genome-wide Repli-seq enrichment of S-phase 2-cell stage embryos and ESCs. This revealed globally organized replication timing patterns recurring within individual cells, with systematic and persistent differences between 2-cell embryos and ESCs (**Fig. 2b-c Supplementary Table 4**). In ESCs, both clusters 2 and 3 replicated late, but in 2-cell embryos their replication timing differed (**Fig. 2c-d**). Cluster 3 replicated comparably early in 2-cell embryos and exhibited high transcriptional induction at the 2-cell stage, while the comparably early replicating ESC cluster 9 was associated with high transcription induction at 4 and 8-cell stage^20^ (**Fig. 2e-f and Extended Data Fig. 2a**). Indeed, previous studies have consistently demonstrated that replication timing is associated with local transcriptional potential and chromatin organization^4,26–28^.

In light of these observations, we sought to investigate the potential influence of maternally inherited epigenetic landscape on the observed differences in replication timing, in addition to transcriptional induction. To accomplish this, we examined the enrichment of epigenetic features using published ChIP-seq, Cut&Run, and ATAC-seq data from Meiosis II (MII) and fully grown oocytes (FGO)^13,14,17,20,29–32^ (**Fig. 2g and Extended Data Fig. 2b-c, Supplementary Table 5**). Interestingly, we identified a pronounced global association between replication timing differences and the H2AUb and H3K27me3 histone marks deposited by Polycomb Repressive Complexes (PRCs)^14,17,29^, which were associated with relatively early replicating regions in the 2-cell embryos compared to ESCs (**Fig. 2g-i and Extended Data Fig. 2b,c, 3a,b**). Together, these analyses revealed that inherited maternal histone marks deposited by PRCs are associated with the temporal organization of DNA replication in 2-cell stage embryos.

### Abundant local parental differences in the replication timing of 2-cell embryos

To explore whether asynchronous DNA replication timing of the parental haplotypes could explain the smaller G1/S deviation in log2 fold signal between 2-cell embryos and ESC (**Fig. 1h-j, Extended Data Fig. 1f,g**), we separated the genomes originating from each of the gametes based on nucleotide variations between the two parental strains. This allowed us to independently visualize and compare the replication timing profiles of the maternal and paternal genome (**Fig. 3a**). Measures of the relative Euclidian distances between Repli-seq values of individual 2-cell and ESC haplotypes as well as hierarchical clustering of these revealed a strikingly distinct DNA replication program in each of the 2-cell haplotypes that was unlike that of ESCs (**Fig. 3b, Extended Data fig. 3c**). Next, we performed k-means clustering of the genome-wide Repli-seq enrichment within the maternal and paternal 2-cell and ESC haplotypes. Based on this, we observed that a large fraction of the genome exhibited consistent earlier replication in the maternal haplotype compared to the paternal haplotype, while the opposite was the case for another large fraction (**Fig. 3c, d, Supplementary Table 6**). These findings in the 2-cell embryo are in contrast to our observations in ESC, where we observed a strong correlation between the replication timing profiles of the two haplotypes, consistent with previous reports (**Fig. 3e**)^9,10^. Notably, separation of paternal haplotypes did not lead to an increase in the low G1/S deviation in log2 fold signal we observed in the zygotes and only partially increased the G1/S deviation of 2-cell embryos, which was still considerably smaller than that of ESCs (**Fig. 1e-f and Extended Data Fig. 3d, e**). This data supports that the lower G1/S deviation observed in zygotes and 2-cell embryos compared to ESCs is not primarily caused by asynchrony between the maternal and paternal genomes. The more uniform progression of replication during early embryogenesis, deviating from the classical temporal pattern, could be the result of actual uniformity or that structures are below our effective resolution.

**Figure 3.**
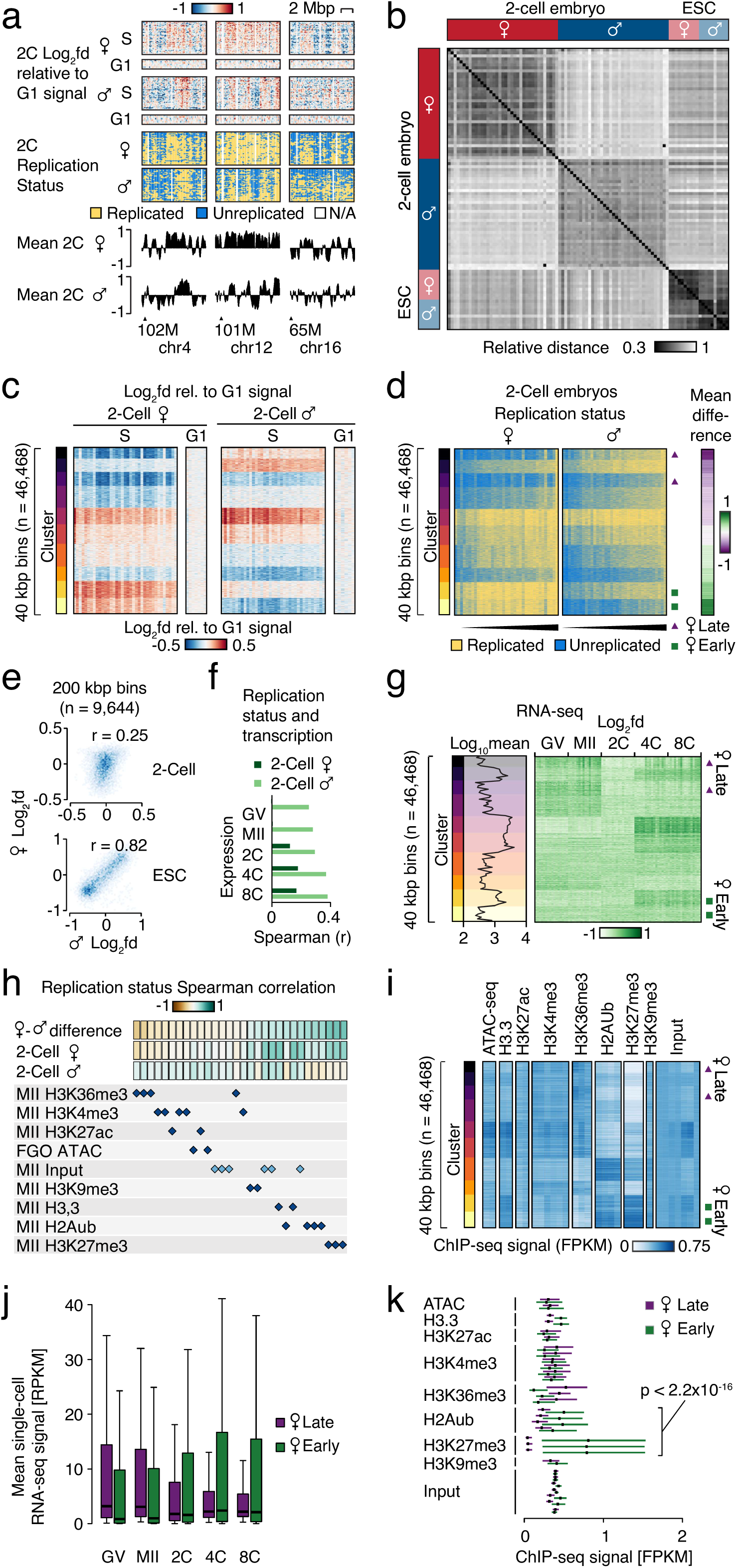
Abundant local parental differences in the replication timing of 2-cell embryos. **a)** Heatmaps showing normalized Repli-seq read densities (upper) and binarized Replication Status (middle) as well as graphs showing mean binarized Replication Status (bottom) of S-phase of the maternal and paternal genome in 2-cell mouse embryos at selected genomic loci. **b)** Distance matrix of showing the relative Euclidian distances between Repli-seq enrichment profiles of maternal and paternal genomes in individual mouse 2-cell (n=37) and ESC samples (n=20). **c)** Heatmaps showing the genome-wide signal enrichment and variation in individual Repli-seq samples from the maternal and paternal genome of S-(n=37) and G1-phase (n=7 Maternal, n=6 Paternal) 2-cell mouse embryos. 40 kbp bins that contained signal from both haplotypes (n = 46,468) were k-means clustered according to the indicated S-phase signal from all individual maternal and paternal samples. Clusters were ordered according to the mean difference between the maternal and paternal signal. **d)** Heatmaps showing the genome-wide binarized Replication Status in individual Repli-seq samples from the maternal and paternal genome of S-phase 2-cell mouse embryos ordered as in (**c**). Rightmost heatmap shows differences in the mean binarized Replication Status between the maternal and paternal genomes. Green squares and purple triangles indicate selected clusters constituting subpopulations of regions with a notably early or late replication of the maternal genome, respectively. **e)** Scatter plots showing the genome-wide relationship between mean normalized S-phase Repli-seq read between the paternal (X-axis) and maternal (Y-axis) genomes for mouse 2-cell embryos (top) or ESCs (bottom) analyzed in 200 kbp bins. R-values indicate Pearson correlation coefficients. **f)** Bar diagrams showing the correlation between mean Replication Status from the maternal and paternal genome of S-phase 2-cell mouse embryos and transcript density from the indicated oocyte and embryo stages. The signal was analyzed in 40 kbp bins and correlation measured using Spearman rank correlation tests. **g)** Heatmaps showing transcript density difference from single mouse oocyte or embryo RNA-seq^19^ Signal was quantified in 40kbp bins ordered as in (**c**) and FPKM-normalized. Green squares: maternal early replicating, purple triangles: maternal late replicating, defined in (**d**). **h)** Tile plot showing the genome-wide correlation of histone mark and chromatin accessibility in mouse oocytes (indicated by lowermost blue rhombs) relative to mean Replication Status from the maternal and paternal genome of S-phase 2-cell mouse embryos as well as the difference between these two. The signal was analyzed in 40 kbp bins and correlation measured using Spearman rank correlation tests. **i)** Genome-wide histone mark and chromatin accessibility signal from mouse oocytes with mean Replication status of the 2-cell embryos and ESC. Signal was quantified in 40kbp bins ordered as in (**c**) and FPKM-normalized. Green squares: maternal early replicating, purple triangles: maternal late replicating, defined in (**d**). **j)** Box plots showing transcript densities at 40 bins within the maternal early and late replicating subpopulations. Whiskers represent the 1.5x interquartile range and the bar is the median value. **k)** Plot of upper and lower quartiles as well as medians (black vertical bars) of histone marks, histone variants, and chromatin accessibility signal from mouse oocytes quantified at 40 kbps within the maternal early and late replicating subpopulations defined in (**d**). P-values obtained by two-sided Mann-Whitney U-tests Benjamini-Hochberg corrected for multiple testing

We next investigated whether genomic regions that harbor high transcriptional induction at the 2-cell stage are replicated early across both parental haplotypes. Indeed, regions of high transcriptional induction are replicated early in both parental haplotypes from 4-cell stage onwards, indicating influence from the major embryonic genome activation^20^ (**Fig. 3f, g and Extended Data fig. 3f**). Given the importance of maternal oocyte derived factors in the development of the embryo, we examined the enrichment of histone marks in relation to replication status differences between the parental haplotypes. Notably, the marks H2AUb and in particularly H3K27me3 deposited by PRCs were enriched in early-replicating maternal regions (**Fig. 3h and i, Extended Data Fig. 4a-d**). Indeed, we observed that the groups of genomic regions with comparably early maternal replication (Defined in **Fig. 3d** and marked by green squares), were characterized by transcriptional induction in the embryo and high maternally inherited H3K27me3 and H2AUb levels (**Fig. 3j and k**). Conversely, regions with comparably late maternal replication (**Fig. 3d**, purple triangles), were characterized by transcriptional downregulation in the embryo and low maternally inherited H3K27me3 and H2AUb levels (**Fig. 3j and k**). Interestingly, the PRC-regulated genomic regions undergoing early replication in the maternal 2-cell genome did not replicate early in ESCs (**Extended Data Fig. 3a, b).** Altogether, this indicates that genome-wide epigenetic diversity across parental genomes is associated with an asynchronous haplotype-specific DNA replication program in the early embryo that is replaced at later stages of embryo development.

### Maternal replication timing is associated with nuclear compartmentalization and epigenetic marks

Remarkably, our mapping of replication timing across whole chromosomes uncovered a compelling pattern. Within a 30-megabase (Mb) segment corresponding to the pericentromeric region in the acrocentric mouse autosomes, there was a distinct overabundance of maternally late replicating regions (**Fig. 4a**). In contrast, no such effect was observed within the paternal chromosomes (**Extended Data Fig. 5a**). This intriguing observation was consistently observed across individual embryos (**Fig. 4b**) and was not a characteristic feature in ESCs (**Extended Data Fig. 5b**). The centromeric regions of the chromosomes in 2-cell embryos congregate at one pole of the nucleus near to the nuclear lamina^33^. Therefore, we hypothesized that Lamin-Associated Domains (LADs) might influence the organization of replication timing at the chromosome level. To test this, we utilized recently published genome-wide LAD data from the maternal and paternal haplotypes of 2-cell embryos^34^. Our analysis revealed a strong association between LADs and replication timing in the 2-cell embryos (**Fig. 4c**), and a particularly pronounced enrichment of LADs in the pericentromeric regions (**Fig. 4d**). Interestingly, both parental haplotypes exhibited a pronounced enrichment of LADs in the late replicating regions (**Fig. 4c and e**). However, the distinct partitioning of late replication in the pericentromeric region was exclusively observed in the maternal genome, while no relationship between the paternal pericentromere and late replication was apparent, despite the accumulation of paternal 2-cell LADs in this region (**Fig. 4d and Extended Data Fig. 5a**). Thus, the pericentromeric DNA is organized within LADs of the 2-cell nucleus and asynchronous parental haplotype replication timing is occurring within this domain.

**Figure 4.**
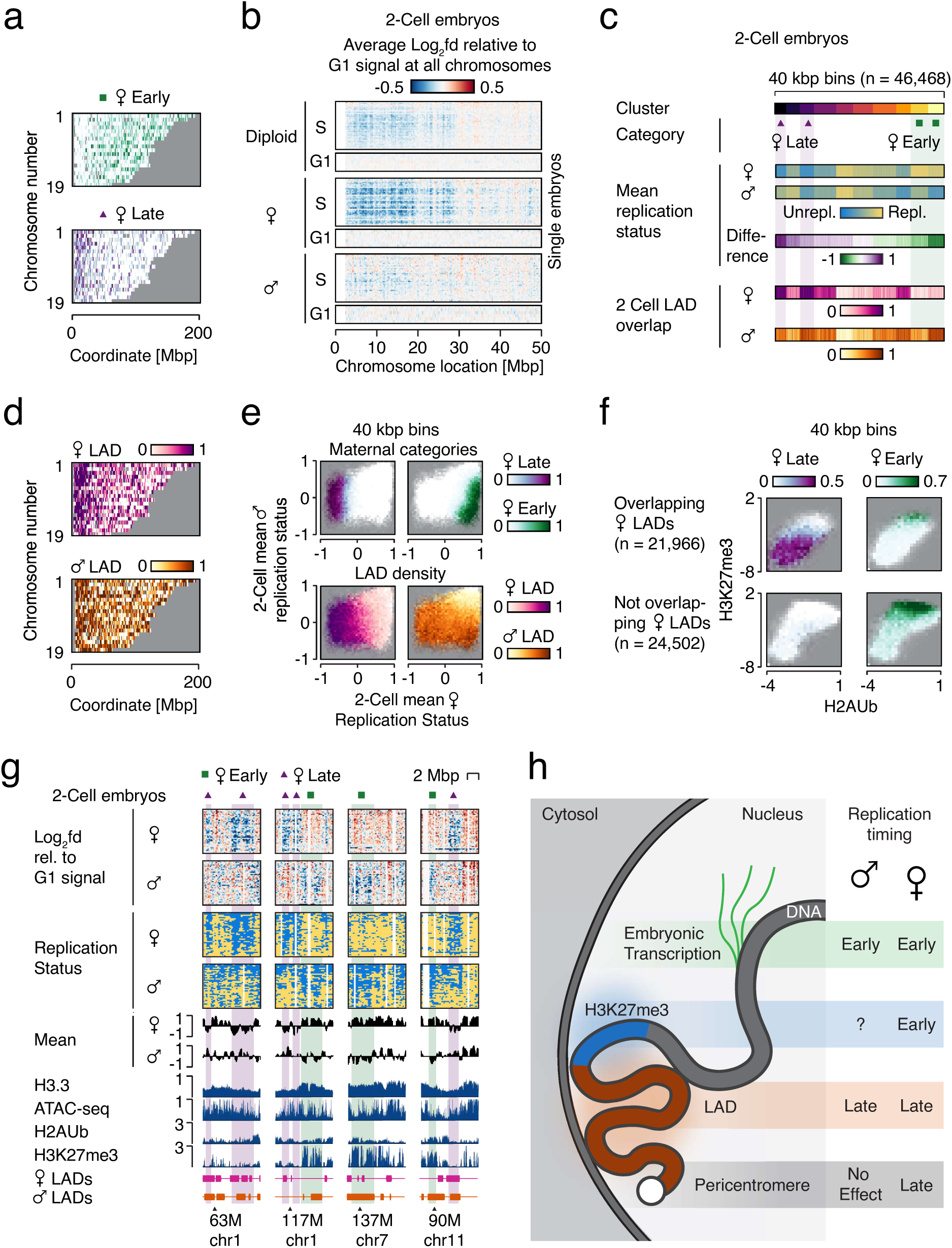
Maternal replication timing is associated with nuclear compartmentalization and epigenetic marks. **a)** Heatmaps showing the density of maternal early (upper) and late (lower) replicating regions, defined by the genome-wide binarized replication status (figure **3d)**. X-axis position correspond to chromosome coordinates, and Y-axes correspond to autosome number. Centromere is positioned at 0 Mbp. **b)** Heatmaps showing the average enrichment at all chromosomes from individual Repli-seq samples from the diploid (top), maternal (middle) and paternal (bottom) genome of S- and G1-phase 2-cell mouse embryos. X-axis position correspond to chromosome coordinates, and Y-axes depict to individual samples. **c)** Heatmaps of genome-wide mean binarized Replication Status and difference (middle) and lamin-associated domains (LAD) density in the maternal and paternal genomes of S-phase 2-cell mouse embryos (bottom) at clusters defined in Figure 3c. **d)** Heatmaps showing the density of maternal (upper) and paternal (lower) LADs in 2-cell mouse embryos. X-axis position correspond to chromosome coordinates, and Y-axes correspond to autosome number. **e)** Heatmaps showing the density of maternal late (upper left) and early (upper right) replicating regions, defined in figure **3d**, as well as the density of maternal (lower left) and paternal (lower right) LADs in 2-cell embryos relative to mean maternal (X-axis) and paternal (Y-axis) binarized Replication Status. **f)** Heatmaps showing the density of maternal late (left) and early (right) replicating regions, defined in figure **3d**, within (top) or outside (bottom) LADs in the maternal genome of 2-cell mouse embryos relative to mean enrichment of the H2AUb (X-axis) and H3K27me3 (Y-axis) histone marks in mouse MII oocytes. **g)** Heatmaps and genome-browser tracks Repli-seq of the maternal and paternal genome in 2-cell S-phase mouse embryos in relation to various features at selected loci. From top: heatmaps of normalized Repli-seq read densities, heatmaps of binarized Replication Status, graphs of mean binarized Replication Status, histone mark, histone variant, and chromatin accessibility in mouse oocytes, maternal and paternal LADs in 2-cell mouse embryos. Regions highlighted in green and orange indicate maternal early and late replication, respectively. **h)** Schematic illustration of identified relationships between maternal and paternal replication timing in 2-cell embryos and key genomic features.

To gain a deeper understanding of the coexistence between maternal late replication of LAD-associated regions and our previous observations of histone marks deposited by PRCs enriched in early-replicating maternal regions, we assessed the combined relationship among PRCs-deposited marks, LADs, and replication timing. Interestingly, our findings revealed that among the histone marks deposited by PRCs, H2AUb but more prominently H3K27me3 were most enriched in the part of the genome that do not overlap with LADs (**Fig. 4f**). In contrast, maternal early replication was predominantly partitioned into non-overlapping LAD regions, which also exhibited high enrichment for marks deposited by PRCs, especially H3K27me3 (**Extended Data Fig. 5c-h**). When we analyzed individual loci we found clear late to early replication timing transitions, which overlapped with boundaries between LADs and H3K27me3 (**Fig. 4g**). These results provide the first demonstration of a large-scale asynchronous haplotype-specific replication program that is influenced by higher-order genome organization, such as LADs, transcription, as well as the inheritance of maternal marks deposited by PRCs (**Fig. 4h**).

## Discussion

The spatiotemporal pattern of DNA replication has been widely recognized as associated with epigenetic marks, transcriptional activity, and the organization of the 3D genome. Across eukaryotes, a DNA replication program exists, which features constant timing of multi-megabase regions ensuring their replication at specific times during the S phase. These regions are flanked by timing transition regions where DNA replication is progressing^35^.

In a departure from the norm, our investigation using single Embryo Repli-seq revealed low G1/S deviation in zygotes, which was only partially restored in the 2-Cell embryo compared to mouse embryonic stem cells (ESCs). We propose that this observation suggests an unconventional DNA replication program in the zygote, characterized by the initiation of a high number of replication origins throughout the genome. The partial recovery of the G1/S signal in 2-cell embryos, compared to ESCs, may indicate a gradual establishment of the replication program as embryonic development unfolds. This would be consistent with the embryonic genome activation and the formation of A/B compartments, which are also established gradually during embryogenesis^36,37^.

Equally fascinating, was the lack of synchrony observed in the 2-cell embryo. Post-translational modifications of histones undergo dynamic changes specific to each parental haplotype following fertilization^38^. We observed a pronounced enrichment of maternally inherited marks deposited by PRCs in the regions that replicate early on the maternal allele. This suggests that the persistence of PRCs-deposited histones from oocytes may play a role in organizing the replication program or be influenced by it.

Furthermore, we made intriguing observations regarding the spatial arrangement of DNA replication in the pericentromeric regions of parental chromosomes, which served to further distinguish the events occurring within the parental haplotypes (**Fig. 4g**). We identified consistent late replication in maternal pericentromeric regions in a large fraction of the studied 2-cell embryos and at most chromosomes (**Fig 4a,b, Extended Data Fig. 5a**), which was also characterized by the absence of histone marks deposited by PRCs. One potential explanation for this observation is the spatial separation of parental genomes within the nucleus, which persists until the 8-cell stage^33^. This separation is achieved by congregation of centromeric regions at one pole of the nucleus, while the long chromosome arms and telomeric regions extend towards the opposite pole^33^. This spatial organization plays a pivotal role in establishing the parental-specific nuclear architecture observed during early development. However, the question of why parental haplotypes display such contrasting replication timing in seemingly similar structural domains, such as LADs should be a question for future investigations.

Given recent reports of heightened replication stress during the cleavage stages of embryogenesis, the exploration of paternal-specific DNA replication stress and stress-induced DNA breaks across the parental haplotypes becomes a matter of great interest. Moreover, the fragile nature of the pericentromere^39–44^ raises intriguing questions about the potential impact of these replication dynamics on the occurrence of gross chromosomal rearrangements, which are relatively common during early embryogenesis^45–47^. Investigations of the role of parental haplotypes in the formation of such rearrangements holds promise for advancing our understanding of embryonic failure and infertility.

In summary, our study unveils a notable link between replication timing and genome organization in the early embryo. Specifically, we have discovered a remarkable excess of late replicating regions in the maternal pericentromeric region when compared to paternal chromosomes. The identification of distinct replication timing patterns between maternal and paternal chromosomes emphasizes the intricate nature of replication regulation and its connection to genome organization. The observed association between replication timing, LADs, and maternal histone marks deposited by PRCs reveals an intriguing interplay between epigenetic modifications, nuclear architecture, and replication dynamics.

## Supporting information

Merged supplementary figures and legends

## Online methods

### mESC Cell Culture

F121-9 mESC line (RRID: CVCL_VC42, a gift from Joost Gribnau) was cultured on 6-well plates previously coated with 0.01% poly-l-ornithine (Sigma P3655) for 1 hour. Subsequently, the plates were further coated with 300pg/ml of laminin (Fisher Scientific 10152421) for a minimum of 1 hour.

The mESCs were maintained in 2i and LIF media as previously described^48^, which was refreshed every 24 hours to ensure optimal growth and viability.

### Sample preparation for single cell RT profiling of mESC cells

Sample preparation of mouse ESCs for single-cell Repliseq was performed following previously established protocols^49,50^. In brief, detached single cells were resuspended in 1 ml of 1% fetal bovine serum (FBS) in phosphate-buffered saline (PBS). Subsequently, the cells were fixed by slowly adding 3 ml of ice-cold 100% ethanol and incubating the mixture at -20°C for 30 minutes.

To ensure accurate cell cycle phase identification, 1×10^6^ cells were washed and resuspended in 700 µl of propidium iodide (PI) staining solution. The PI staining solution comprised of 350 µl of 1 mg/ml PI, 70 µl of 10 mg/ml RNase A, and 7 ml of 1% FBS-PBS.

Flow cytometric sorting was performed using a Sony SH800 sorter to isolate single cells in the G1 or S phase, based on their PI staining profile. The sorted single cells were collected directly into individual wells of a 96-well plate. Each well contained 6 µl of single-cell lysis and fragmentation buffer, comprising of 3 µl of proteinase K, 50 µl of 10x Single Cell Lysis and Fragmentation Buffer (Sigma; L1043), and 447 µl of autoclaved double-distilled water (ddH2O).

Following collection, the plate was briefly centrifuged and subsequently stored at -80°C until further processing for single-cell sequencing.

### Spermatozoa cryopreservation

For cryopreservation of mouse sperm, a modified version of a previously established protocol was utilized^51^.

In brief, ten 0.25 mL clear plastic straws (Minitube GmbH; 13407/0010) were labeled for each individual mouse. A proven fertile male mouse, aged between 3 to 8 months, was sacrificed following one week of solitary housing. The cauda epididymis and vas deferens were dissected and placed in 1 mL of Dulbecco’s phosphate-buffered saline (DPBS). Under a dissection microscope, adipose and vascular tissues were carefully removed. Both cauda epididymides were then transferred to a 35mm dish containing a 120 µl droplet of gCPA sperm cryopreservation media (in-house) covered with NidOil (Nidacon; NO-100).

Using a pair of watchmaker scissors, the epididymides were cut several times to release the sperm and facilitate its separation from the tissue. The dish was placed in a CO2 incubator at 37°C with 5% CO2. To aid sperm dispersion and release, the dish was gently swirled once every minute for a total of 3 minutes.

Ten 10 µl aliquots of the sperm suspension were taken from the dish and deposited onto the lid of a 35mm culture dish. Clear plastic straws were prepared for freezing by drawing up HTF medium (developed in-house) into the straw using a syringe, followed by a small amount of air, and then the 10 µl aliquot of sperm. Additional air was introduced into the straw before sealing it with a metal ball (Minitube GmbH; 13400/9970).

All ten straws were then placed in the gaseous phase of liquid nitrogen for a duration of 10 minutes. Subsequently, the straws were slowly and carefully immersed into the liquid phase of nitrogen before being transferred to a -150°C freezer or long-term liquid nitrogen storage for preservation.

### *In Vitro* fertilization of mouse oocytes

*In vitro* fertilization of mouse oocytes was performed as previously described but with some modifications^51^. Briefly, a single straw of cryopreserved sperm from CAST/EiJ mice was thawed for 10 minutes in a water bath set to 37°C. After thawing, the sperm was dispersed into 90 µl of TYH+MBCD (in house) sperm pre-incubation medium in a 35mm dish coated with NidOil (Nidacon; NO-100). The dish was equilibrated at 37°C for 20 minutes in a 5% CO_2_ in air environment.

Simultaneously, a 30mm fertilization dish was prepared by adding 90 µl droplet of 2 mM reduced glutathione (Sigma, G4251) in HTF (in house) and covering it with NidOil. The dish was equilibrated for at least 30 minutes at 37°C in a 5% CO2 in air environment.

Female C57/BL6N mice were superovulated by intraperitoneal injection of 7.5 IU of hCG (Sigma; CG5) and 7.5 IU of PMSG (Prospec; HOR-272) 48 hours later. After 15 hours from the second injection, the mice were euthanized, and the oviducts were dissected and placed into the warm NidOil of the fertilization dish. Cumulus-enclosed oocytes were transferred from the ampoule into the fertilization droplet.

Following a 30-minute pre-incubation, 10 µl of the most motile sperm was extracted from the pre-incubation droplet and added to the fertilization dish containing the cumulus-enclosed oocytes. The dish was then incubated for three hours.

Meanwhile, a wash dish was prepared by creating four droplets of 90 µl of Advanced KSOM embryo media (Sigma; MR-101) in a 60mm dish. The dish was covered with NidOil and equilibrated for 30 minutes at 37°C in a 5% CO2 environment.

Subsequently, the fertilized oocytes were washed through the four wash droplets to remove the cumulus cells and sperm. The oocytes were then incubated until they were ready for Single-Embryo Repli-Seq.

### EdU labelling of single mouse embryos

To determine the duration of the first and second S phase, we conducted pulse labeling of single 2-cell stage embryos using the Click-iT EdU Cell Proliferation Kit for Imaging (ThermoFisher Scientific; C10340). The incorporation of EdU was assessed through microscopy. The experimental procedure was as follows:

Embryos were obtained at the zygote (no prior treatment) or the 2-cell stage following release from Nocodazole synchronization. At hourly intervals, individual zygotes or 2-cell embryos were transferred to pre-equilibrated droplets of Advanced KSOM embryo media (Sigma; MR-101) containing 500 µM EdU. The droplets were maintained at 37°C in a 5% CO2 environment. After a 30-minute incubation period, the embryos were washed with Advanced KSOM media and fixed in 4% paraformaldehyde (PFA) for 15 minutes at room temperature. Subsequently, the embryos were transferred to a wash buffer consisting of 5% BSA in PBS and incubated on a rocking stage at 4°C for 48 hours. To permeabilize the embryos, they were treated with 0.5% Triton X-100 in PBS for 20 minutes at room temperature, followed by rinsing in PBST (0.1% Tween 20 in PBS) for 20 minutes at room temperature. The embryos were then subjected to the Click-iT EdU cell proliferation reaction cocktail, according to the manufacturer’s instructions. They were incubated in the reaction cocktail for 1 hour. After the incubation, the embryos were rinsed in a wash buffer containing 20 µg/ml DAPI (ThermoFisher Scientific; D1306) for 30 minutes.

For imaging, embryos from the same time point were mounted using SlowFade Diamond Antifade Mountant (ThermoFisher Scientific; S36967) between two No.1 coverslip. Confocal microscopy imaging was performed using an LSM900 confocal laser scanning microscope (Zeiss) equipped with a 40x Plan-Apochromat objective. The Alexa 647 fluorophore was excited using a 639-nm laser, and DAPI was excited using a 405-nm laser. Image processing was performed using Zen software and ImageJ^52^ software for quantification of corrected total fluorescence.

### Sample preparation for RT profiling of single mouse embryos (Single-Embryo Repli-Seq)

Zygotes collected after IVF were categorized into G1 or S phase. G1 phase was determined by two pronuclei detection, while S phase was determined by timing established using prior EdU incorporation experiments. Zona pellucida and polar bodies were removed using a dissection microscope, and naked zygotes were transferred to a low-adhesion 96-well PCR plate. Each well contained 6 µl of single-cell lysis and fragmentation buffer (3 µl proteinase K, 50 µl 10x Single Cell Lysis and Fragmentation Buffer (Sigma; L1043), and 447 µl ddH2O). The plate was centrifuged and stored at -80°C for single-cell sequencing.

For 2-cell embryos, fifteen hours after combining sperm with cumulus-enclosed oocytes, the resulting zygotes were transferred using a mouth pipette into a 90 µl droplet of Advanced KSOM embryo media (Sigma; MR-101) containing 40 nM Nocodazole. The droplet was placed in a 30mm dish covered with NidOil (Nidacon; NO-100). This step synchronized the zygotes in the G2/M phase from which they were released after 4 hours.

Following the synchronization period, the zygotes were washed through seven droplets of Advanced KSOM embryo media, each covered with NidOil. These droplets were pre-equilibrated for 30 minutes at 37°C in a 5% CO2 environment.

Under normal circumstances, the zygotes divide within 1 hour after release from the block. At this point, G1 phase 2-cell stage embryos were promptly processed by removing the zona pellucida and polar bodies (if present) using a dissection microscope. The resulting naked embryos were then transferred to a low-adhesion 96-well PCR plate. Each well contained 6 µl of single-cell lysis and fragmentation buffer. The plate was briefly centrifuged and subsequently stored at -80°C until preparation for single-cell sequencing.

For the processing of single S-phase embryos, the same procedure was followed. However, accurate selection of S-phase embryos was based on prior EdU labeling experiments, which determined the time of entry and exit from the S phase after release from Nocodazole synchronization. Specifically, early S-phase cells were selected 3 hours after release, mid-S-phase cells were selected 5 hours after release, and late-S-phase cells were selected 6 hours after release.

### Sample preparation and sequencing for single embryo RT profiling

After sample preparation, single embryos or single mouse embryonic stem cells (mESCs) were processed for single-cell sequencing following a previously published single-cell Reli-Seq protocol^49,50^. After whole-genome amplification, the samples were quantified using the Qubit dsDNA HS assay kit (ThermoFisher; Q32851). The quantification of the samples was further confirmed by checking the product size using the Agilent Tapestation 2200 High Sensitivity system (Agilent; 5067). For library preparation, we utilized the KAPA Hyper Prep Library Preparation Kit (Roche; KK8502) in combination with IDT for Illumina TruSeq DNA UD Indexes (Illumina; 20023784). The end repair and A-tailing steps were performed using 40 ng of whole-genome amplified DNA. The reaction volume was scaled down to 1/5th of the recommended reaction volume and vortexed and briefly spun down before incubating on a thermocycler at 20°C for 30 minutes, followed by 65°C for 30 minutes, and then held at 4°C. Adaptor ligation was performed in the same plate as the end repair and A-tailing reaction. The reaction mixture consisted of 12 µl end repair and A-tailing reaction product, 0.5 µl IDT for Illumina TruSeq DNA UD Indexes, 1.5 µl PCR-grade water, 6 µl ligation buffer, and 2 µl DNA ligase. The reaction mixture was vortexed, spun down, and incubated at 20°C for 15 minutes. Post-adaptor ligation cleanup was performed using 0.8X Agencourt AMPure XP beads (Beckman Coulter; A63881), following the instructions provided with the KAPA Hyper Prep Kit. Library amplification was carried out by scaling down the recommended manufacturer’s protocol to 1/5^th^ and using 11 µl of the bead-cleaned post-ligation product. Amplification was performed using a thermocycler with an initial denaturation step of 98°C for 45 seconds, followed by 5 PCR cycles at 98°C for 15 seconds, 60°C for 30 seconds, and 72°C for 30 seconds. After the final cycle, an extension step was included, and the samples were incubated at 72°C for 1 minute, followed by holding at 4°C. The samples were then cleaned according to the KAPA Hyper Prep Kit post-amplification cleanup instructions using 0.9X Agencourt AMPure XP beads. The final libraries were quantified using the Qubit dsDNA HS kit and assessed for library size using the Agilent Tapestation 2200 High Sensitivity system. The libraries were sequenced on a NextSeq550 platform using a 150-bp single-end read format.

### Single-Embryo and single mESC Repli-Seq data analysis

Our single embryo Repli-Seq data analysis, which was used to analyse all embryos and single mESCs, was based on two previously published protocols^49,50,53^. We followed the analysis pipeline described by Miura et al. (2019) and Miura et al. (2020) up to the generation of log2 fold replication timing (RT) scores, with some modifications. The binarization of Replication Status followed the piecewise copy fit and mixture modelling approach outlined by Dileep and Gilbert (2018) with adaptations for our analysis. This relied on the R-packages Zoo, Fbasics, Mixtools, and Copynumber^54^.

Fastq files for G1 and S phase were checked for quality using FastQC^55,56^ and trimmed for Illumina index and Seqplex adaptor sequences using Cutadapt^57^, as described in previous studies^49,50^. For haplotype analysis, sequencing reads were mapped to a modified mm10 reference genome that included N-masked positions for single hybrid strains. The N masked genome was created using the genome preparation command of the SNPsplit package^58^, which incorporates N-masking into the mm10 reference for the CAST/EiJ genome from the Mouse genome project V7 vcf file^59–61^. Subsequently, this new mm10 N-masked genome was then used to map the trimmed sequencing reads using bowtie2^62^ for parsed haplotype analysis, alternatively unparsed haplotype analysis was performed using the original mm10 reference genome and duplicate reads were marked with Picard (http://broadinstitute.github.io/picard).

To divide the reads into maternal and paternal haplotypes, we used the allele-specific alignment sorter SNPsplit^58^, which utilized a list of known SNP positions between C57BL/6 and CAST/EiJ mice. Quality control of parsed BAM files was confirmed using SAMStat^63,64^, and typically, we obtained >50% reads with >30 MAPQ. To calculate log2 fold replication timing scores, a G1 reference file was generated from more than three G1 control cells with a normal karyotype. The karyotype was confirmed using the hidden Markov model of the findCNVs command in the Aneufinder package^65^, as previously described^49,50^. In our analysis of mESC, we made the decision to exclude chromosome 3 and chromosome 8 due to aneuploidy.

Computing of RT scores were performed by counting reads in sliding windows of 200 kb at 40 kb intervals normalized to the G1 control according to the standard scRepliseq analysis or haplotype resolved scRepliseq analysis previously published protocol for unphased and phased haplotype analysis, respectively^50^. Log2 fold RT scores were converted to binary RT scores using a previously outlined protocol, with binary values determined using the skew approach when the component means from the mixture modelling were <0.25^53^.

The Replication Score for each sample was obtained by calculating the fraction of the genome that had been replicated based on the binary scores for each sample, ignoring bins from masked parts of the genome. A matrix assembly was generated containing Normalized Repli-seq enrichment and binarized Replication Status for each cell. Data aggregation for this was done using the merge command in R (v. 4.1.2)^66^ and IDs for each bin consisting of chromosome name and start coordinate for all datasets. This was performed separately for three pairwise comparisons, published vs. our ESC data, ESC vs. 2-cell data, and maternal vs. paternal 2-cell data, resulting in output that only included the relevant datasets and covered parts of the genome where data were present in all input samples (**Supplementary Tables 1,4,6**). Subsequent analysis was performed in EaSeq (see below)^67^ after importing bedgraphs-files of log2-fold enrichment comparisons between S-phase and G1-phase samples as well as bedgraphs of Binarized Replication Status as ‘datasets’.

### Genomic data sources and quality control

Previously published Repli-seq data from single ESCs^53^ were downloaded from the NCBI GEO database^68^, accession GSE102077. Lamin-associated domains in maternal and paternal 2-cell embryos were identified in^69^ and bed-files with allele-specific coordinates were downloaded from the NCBI GEO database^68^, accession GSE112551. Refseq genome annotations^70^ for mm10 were downloaded from the UCSC table browser^71^. Read qualities were analyzed using FastQC^56^, fastqScreen (v. 0.11.4)^72^, and MultiQC (v. 1.7)^73^.

### ChIP-seq, Cut&Run, RNA-seq, and ATAC-seq data sourcing and processing

ChIP-seq of histone marks and corresponding input from MII oocytes were published in^74–78^, Cut&Run of the histone variant H3.3 in MII oocytes published in^79^, and ATAC-seq from fully grown oocytes published^80^. A complete list can be found in **Supplementary Table 3**. Single oocyte / embryo RNA-seq was previously published^78^ and was processed as described in^81^. ChIP-seq, Cut&Run, and ATAC-seq reads were analyzed using FastQC and trimmed using Trimgalore^82^. Reads were mapped using bowtie2^62^, and handled using Samtools^83^. Blacklisted regions^84^ were obtained from https://github.com/Boyle-Lab/Blacklist/archive/v2.0.zip and removed using bedtools intersect -v^85^. Subsequent analysis was performed in EaSeq (see below)^67^ after importing mapped reads from Bam-files as ‘datasets’. For ChIP-seq, Cut&Run and ATAC-seq this was done using default settings, including deduplication. For RNA-seq removal of duplicates was disabled.

### Repli-seq analysis and visualization

Analysis and visualization was done using EaSeq (v. 1.12)^67^, Microsoft Excel 2016 for simple bar diagrams and line charts, and R (v. 4.1.2)^66^ for Boxplots and interquartile range plots as well as the beeswarm package (v.0.4.0, https://github.com/aroneklund/beeswarm) for Bee-swarm plots. In EaSeq, Genome browser tracks and annotations were made using the EaSeq plot tools ‘FillTrack’, ‘Annot.’ and ‘RegionMap’ for LADs (available in the beta-testing panel). Scatter plots and heatmaps of ChIP-seq, Cut&Run, ATAC-seq, and RNA-seq values were made using the EaSeq tools ‘Scatter’ and ‘ParMap’. Assembled matrices (**Supplementary Tables 4-6**) and LAD coordinates we imported as ‘Regionsets’ in EaSeq. Heatmaps of single-embryo log2-fold enrichment comparisons between S-phase and G1-phase samples as well as heatmaps of Binarized Replication Status at individual loci (as well as averages from multiple loci) were generated using the ‘Multi Track’ tool available in the ‘Beta-testing’ panel of EaSeq. ChIP-seq, Cut&Run, and ATAC-seq Signal intensities at the bins in the matrix assemblies were quantified using the EaSeq tool ‘Quantify’, using default normalization (Fragments Per Kilobase per Million reads – FPKM) for scatter plots and clustered heatmaps. Similarly, RNA-seq read densities were quantified using the EaSeq tool ‘Quantify’ but with normalization disabled to obtain matrices of raw read counts. Following this, a pseudoread was added to RNA-seq read counts, and all quantified RNA-seq counts in a matrix assembly were quantile normalized using the tool ‘Normalize’. These normalized values were used to calculate means for each condition as well as log2 fold difference from the mean using the EaSeq tool ‘Calculate’. Heatmaps of feature density e.g. ChIP-seq, RNA-seq signal, or LAD-overlap were generated using the tool ‘Z-scatter’. Similarly, plots of features at autosomes were generated using this tool and the autosome number for the Y-axis and start coordinate of each bin for the X-axis. Overlap with LADs for each matrix assembly was determined using the EaSeq tool ‘Coloc’. Subpopulations were obtained using the ‘Gate’ tool.

### Repliseq clustering

Hierarchical clustering was performed in R (v. 4.1.2)^66^ on scaled and transformed (R-commands: scale() and t(), default settings) log2-fold Repli-seq enrichment using the command dist(method = ’euclidean’) to calculate a distance matrix, and a dendrogram was plotted after hierarchically clustering the distance matrix using the command hclust(method = ’ward.D2’). k-means-clustering was performed in EaSeq, using the tool ‘ClusterP with all additional normalization disabled. Clustering was performed individually for comparisons between 2-cell and ESC data and 2-cell maternal and paternal data.

### Statistics

All statistics was performed as described in legends using R (v. 4.1.2)^66^.

## Code availability

Scripts for diploid and haplotype-resolved Repli-seq processing were acquired from https://github.com/kuzobuta/scRepliseq-Pipeline and https://github.com/kuzobuta/scRepliseq-Pipeline/tree/master/scripts/haplotype-resolved-analysis. EaSeq and source code is available at http://easeq.net

## Data availability

**FOR REVIEWERS (will be deleted from the manuscript upon acceptance):**

**To review GEO accession GSE237400 (First submission data):**

Reviewer access to the unpublished Repli-seq data for the first submission of this study can be obtained using the link https://www.ncbi.nlm.nih.gov/geo/query/acc.cgi?acc=GSE237400 and by entering the token abyfqaiwjrwnrob into the box.

The Repli-seq data generated for this publication have been deposited in NCBI’s Gene Expression Omnibus^68^ and are accessible through GEO Series accession number GSE237400 (https://www.ncbi.nlm.nih.gov/geo/query/acc.cgi?acc=GSE237400).

## Acknowledgements

We thank the members of the Hoffmann lab and Center for Chromosome Stability for support and critical suggestions throughout this work. We thank Ricardo Alonso Laguna Barraza, Rodrigo Dos A Garcia and Ioana Maria Zah of the Core Facility of Transgenic Mice, The University of Copenhagen for their technical support. We acknowledge the Core Facility for Integrated Microscopy, Faculty of Health and Medical Sciences, University of Copenhagen. We would like to express our gratitude to David Gilbert and Vishnu Dileep for their valuable guidance on single-cell Repli-Seq methodology. Additionally, we appreciate the insightful discussions and assistance provided by Ichiro Hiratani and Hisashi Miura during the initial phase of establishing the single-cell Repli-Seq methodology

## Author contributions

J.A.H., E.R.H. and M.L. conceived the study and designed experiments. J.A.H. and J.M.G. planned and handled mouse work, IVF, and embryo collection. Embryo culture, labeling, imaging, and experiments was carried out by J.A.H. Library preparation was conducted by J.A.H. Data analysis and processing was conducted by J.A.H., A.H., and M.L. J.A.H., J.A.D, E.R.H. and M.L. discussed data and provided critical input. J.A.D., E.R.H. and M.L. prepared the manuscript. All authors read and commented on the manuscript.

