## Supplementary material for "Sex-Specific DNA-Replication In The Early Mammalian Embryo": Merged supplementary figures and legends

Halliwell et al., Extended Data Figure 1

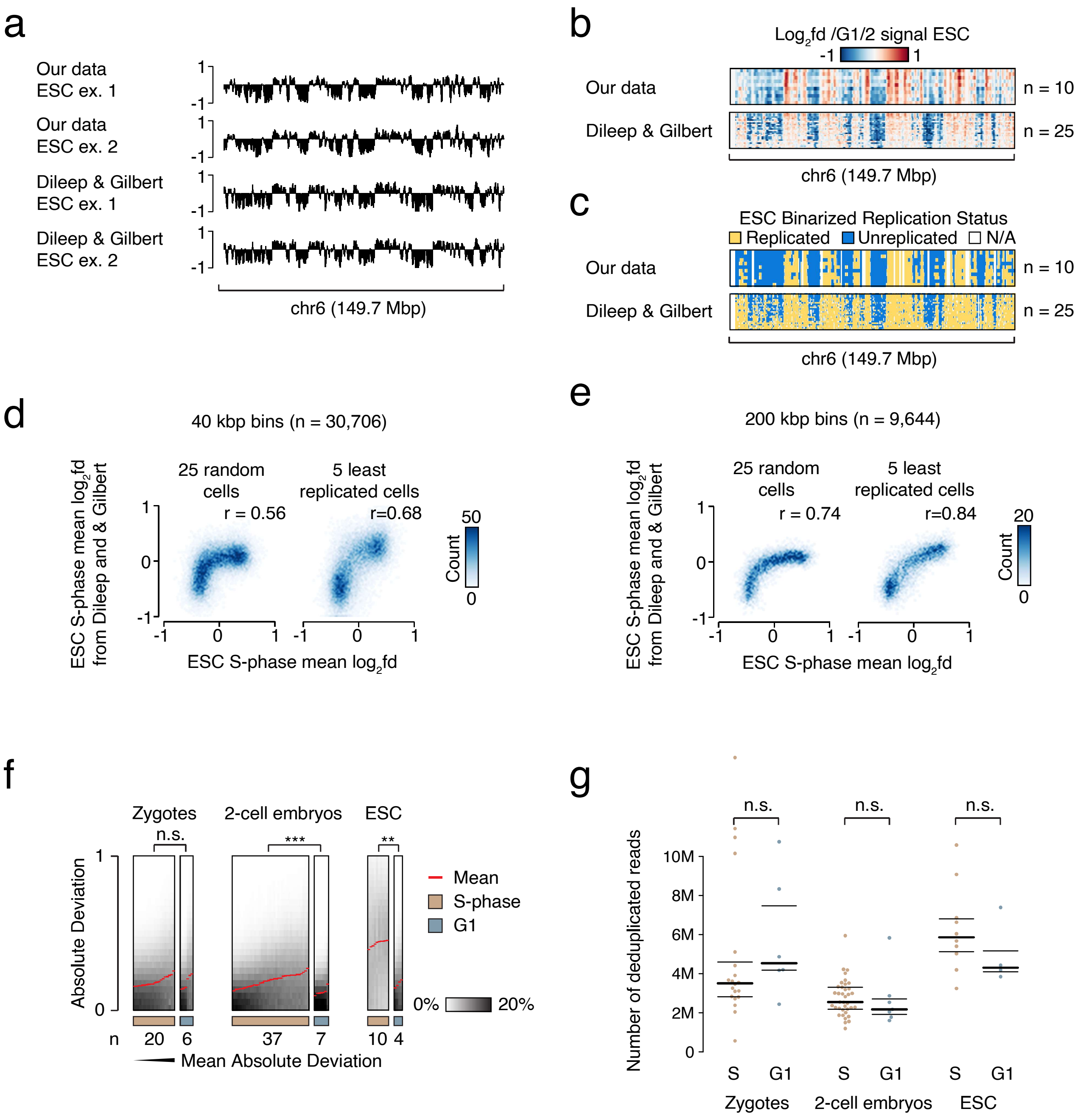


**Single-embryo Repli-seq profiles aligns with previously published sc-Repli-seq profiles.**
**a**) Graphs showing normalized Repli-seq read densities in two S-phase mouse ESCs compared to similar profiles from two published mouse ESCs[9] at chromosome 6. Values are log2-fold differences relative to the G1-phase signal.
**b**) Heatmaps showing individual normalized Repli-seq read densities in S-phase mouse ESCs (n=10) compared to similar profiles from published mouse ESCs[9] (n=25) at chromosome 6. Values are log2-fold differences relative to the G1-phase signal for each cell type.
**c**) Heatmaps showing individual binarized Replication Status of S-phase mouse ESCs (n=10) compared to similar profiles from published mouse ESCs[9] (n=25) at chromosome 6. Binarized values were derived from Repli-seq data normalized to the G1-phase signal for each source.
**d, e**) Scatter plots showing the relationship between mean normalized of S-phase mouse ESCs (X-axis) compared to similar profiles from published mouse ESCs (Y-axis) quantified in 40 kbp windows (**d**) or 200 kbp windows (**e**). Left side plots are based on 25 random published cells[9], whereas the rightmost plot are based on the five of these cells with the lowest binarized Replication Score. R-values indicate Pearson correlation coefficients.
**f**) Heatmaps of our absolute genome-wide deviation in individual Repli-seq S- and G1- phase samples ranked according to the mean absolute deviation (red curves). P-values obtained by two-sided Mann-Whitney U-tests Benjamini-Hochberg corrected for multiple testing. ***: adjusted p-value < 0.001; **: adjusted p-value < 0.01; n.s.: not significant.
**g**) Beeswarm plots of the number of deduplicated mapped reads in each Repli-seq sample. Black bars indicate the 25, 50, and 75 percentiles. P-values obtained by two-sided Welch Two-sample t-tests showing no significance (no correction for multiple testing).

### Halliwell et al., Extended Data Figure 2

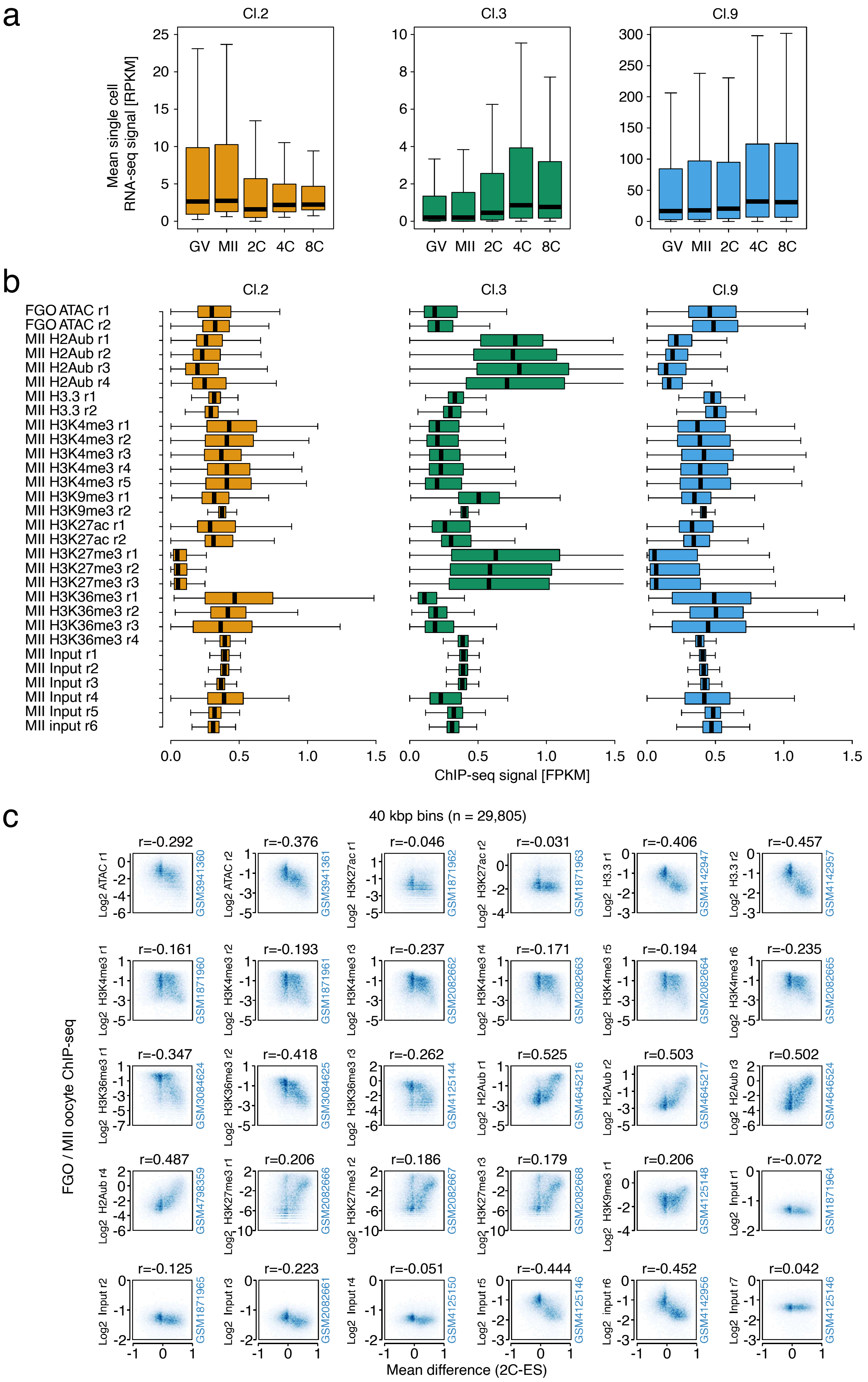

### Halliwell et al., Extended Data Figure 2 Legends

**ESC and 2-cell replication timing disparities correlate with both transcriptional activity and various epigenetic marks.**

**a**) Boxplots showing the transcript density in single mouse oocyte or embryo RNA-seq quantified in 40 kbp bins within three selected clusters (clusters 2, 3, and 9) from Figure 2c. Horizontal bands show medians, bars the interquartile range, and whiskers data points within 1.5x the interquartile range of the lowest and highest quartiles. RPKM: Reads per kilobasepair per million.

**b**) Boxplots showing the density of histone marks, histone variants, and chromatin accessibility in mouse oocytes quantified in 40 kbp bins within three selected clusters (clusters 2, 3, and 9) from Figure 2b. Horizontal bands show medians, bars the interquartile range, and whiskers data points within 1.5x the interquartile range of the lowest and highest quartiles. FPKM: Fragments per kilobasepair per million.

**c**) Scatter plots showing the genome-wide relationship of the mean binarized Replication Status difference between mouse 2-cell embryos and ESCs (X-axis) compared to mouse oocyte histone marks, histone variants, and chromatin accessibility (Y-axis). Values were quantified in 40 kbp windows and r-values indicate Pearson correlation coefficients. Blue vertical numbers are GEO accession numbers for individual datasets.

Halliwell et al., Extended Data Figure 3

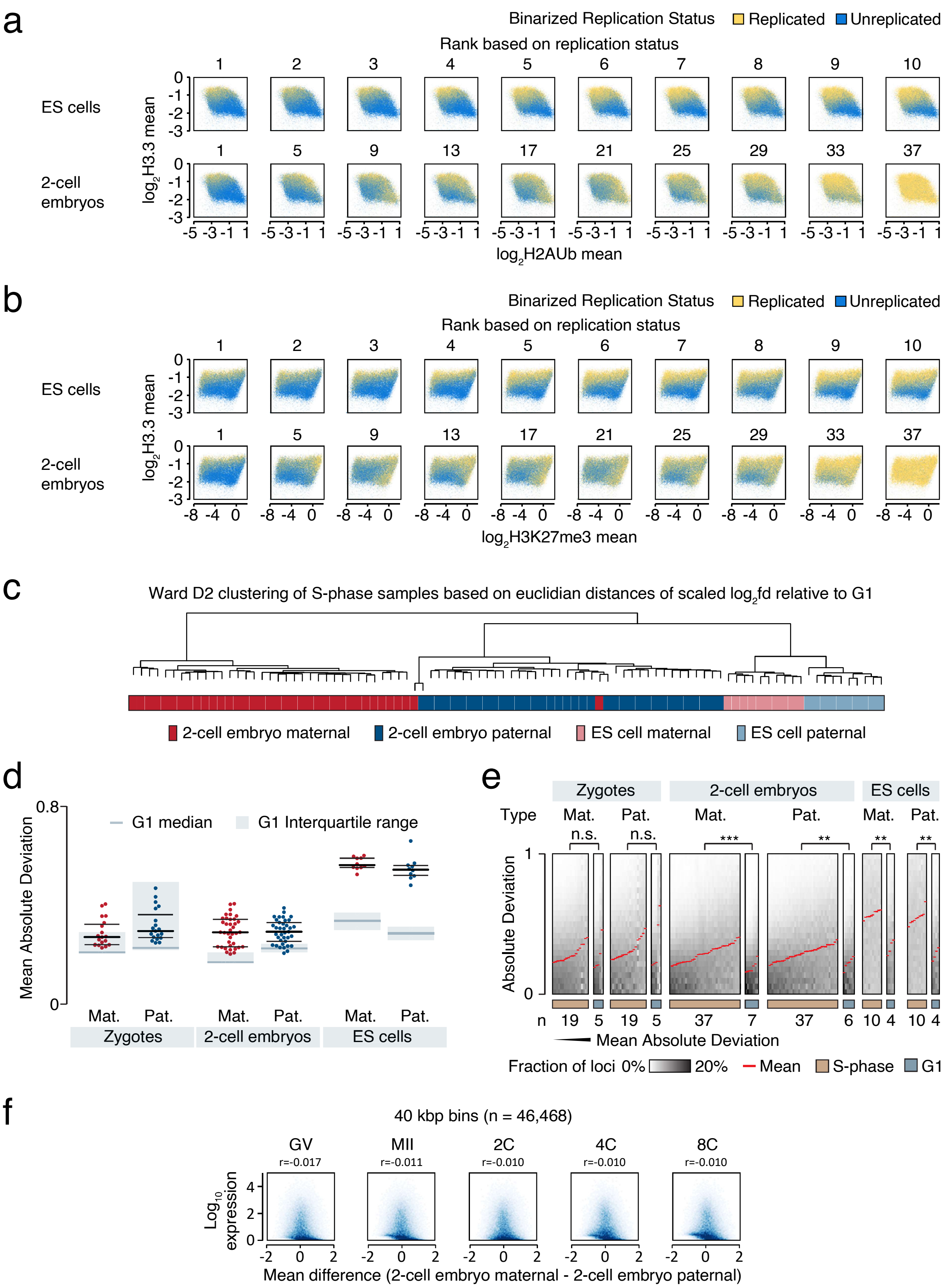


**Lower G1/S deviation in early embryos compared to ESCs isn't primarily due to genome asynchrony.**

**a, b**) Heatmaps showing the genome-wide relationships between mean binarized Replication Status in individual ESCs (top) and mouse 2-cell embryos (bottom) relative to (**a**) mean enrichment of the H2AUb histone marks (X-axis) and H3.3 histone variant (Y-axis), or (**b**) relative to mean enrichment of the H3K27me3 histone mark (X-axis) and H3.3 histone variant (Y-axis) in mouse MII oocytes. Values were quantified in 40 kbp windows. The horizontal order of each heatmap (and header) reflects the ranking based on the replication status for each sample. For 2-cell embryos, only every fifth sample was shown.

**c**) Dendrogram based on hierarchical clustering of Euclidian distances between normalized Repli-seq enrichment profiles of maternal and paternal genomes in individual mouse 2-cell and ESCs S-phase samples. Values were log2-fold differences relative to the G1-phase signal and scaled to have equal mean and standard deviations prior to Wards D2 clustering.

(*Continues on next page*)

**d**) Beeswarm plots of the mean absolute genome-wide deviation within the maternal and paternal genomes of S-phase 2-cell mouse embryos in each Repli-seq sample. The background deviation of G1 controls from each sample type is indicated with grey boxes and wide bars showing the interquartile range and median, respectively. Black bars indicate the 25, 50, and 75 percentiles of the S-phase populations.
**e**) Heatmaps of the absolute genome-wide deviation within the maternal and paternal genomes of S-phase 2-cell mouse embryos in individual Repli-seq S- and G1- phase samples ranked according to the mean absolute deviation (red curves). P-values obtained by two-sided Mann-Whitney U-tests Benjamini-Hochberg corrected for multiple testing. ***: adjusted p-value < 0.001; **: adjusted p-value < 0.01; n.s.: not significant.
**f**) Scatter plots showing the genome-wide relationship of the mean binarized Replication Status difference between maternal and paternal genomes of S-phase 2-cell mouse embryos (X-axis) compared to mean RNA-seq transcript density from single mouse oocyte or embryo (Y-axis). Values were quantified in 40 kbp windows and r-values indicate Pearson correlation coefficients.

Halliwell et al., Extended Data Figure 4

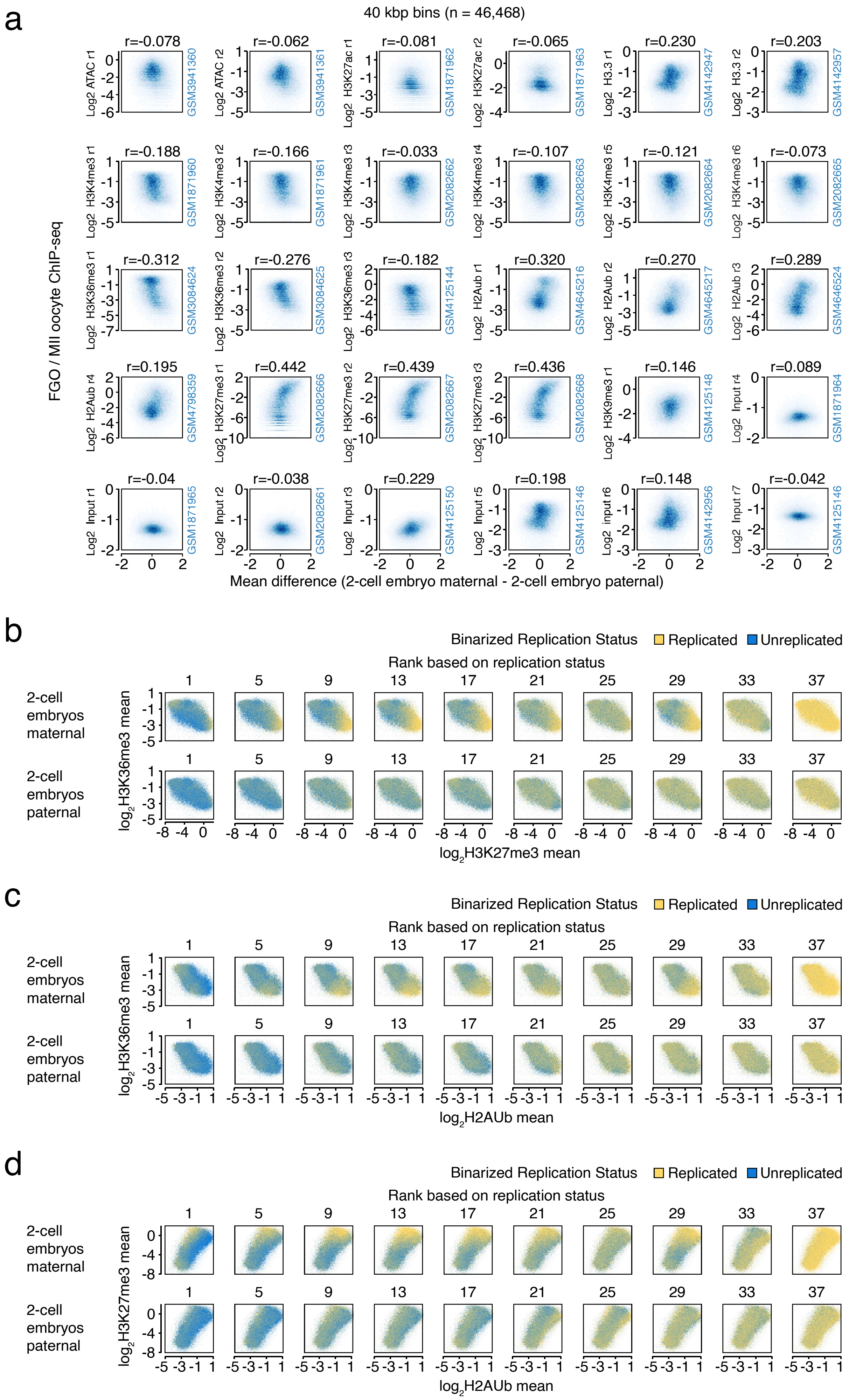

**Early-replicating maternal regions showed enrichment of epigenetic marks deposited by PRCs.**

**a**) Scatter plots showing the genome-wide relationship of the mean binarized Replication Status difference between maternal and paternal genomes of 2-cell mouse S-phase embryos (X-axis) compared to mouse oocyte histone marks, histone variants, and chromatin accessibility (Y-axis). Values were quantified in 40 kbp windows and r-values indicate Pearson correlation coefficients. Blue vertical numbers are GEO accession numbers for individual datasets.

**b-d**) Heatmaps showing the genome-wide relationships between mean binarized Replication Status in individual maternal (top) and paternal (bottom) genomes of 2-cell mouse S-phase embryos and selected histone marks in mouse MII oocytes (X- and Y-axes). Axes represent: (**b**) mean enrichment of the H3K27me3 (X-axis) and H3K36me3 (Y-axis) histone marks, (**c**) mean enrichment of the H2AUb (X-axis) and H3K36me3 (Y-axis) histone marks, and (**d**) mean enrichment of the H2AUb (X-axis) and H3K27me3 (Y-axis) histone marks. Values were quantified in 40 kbp windows. The horizontal order of each heatmap (and header) reflects the ranking based on the sum of the binarized values for each sample. Only every fifth sample is shown.

### Halliwell et al., Extended Data Figure 5

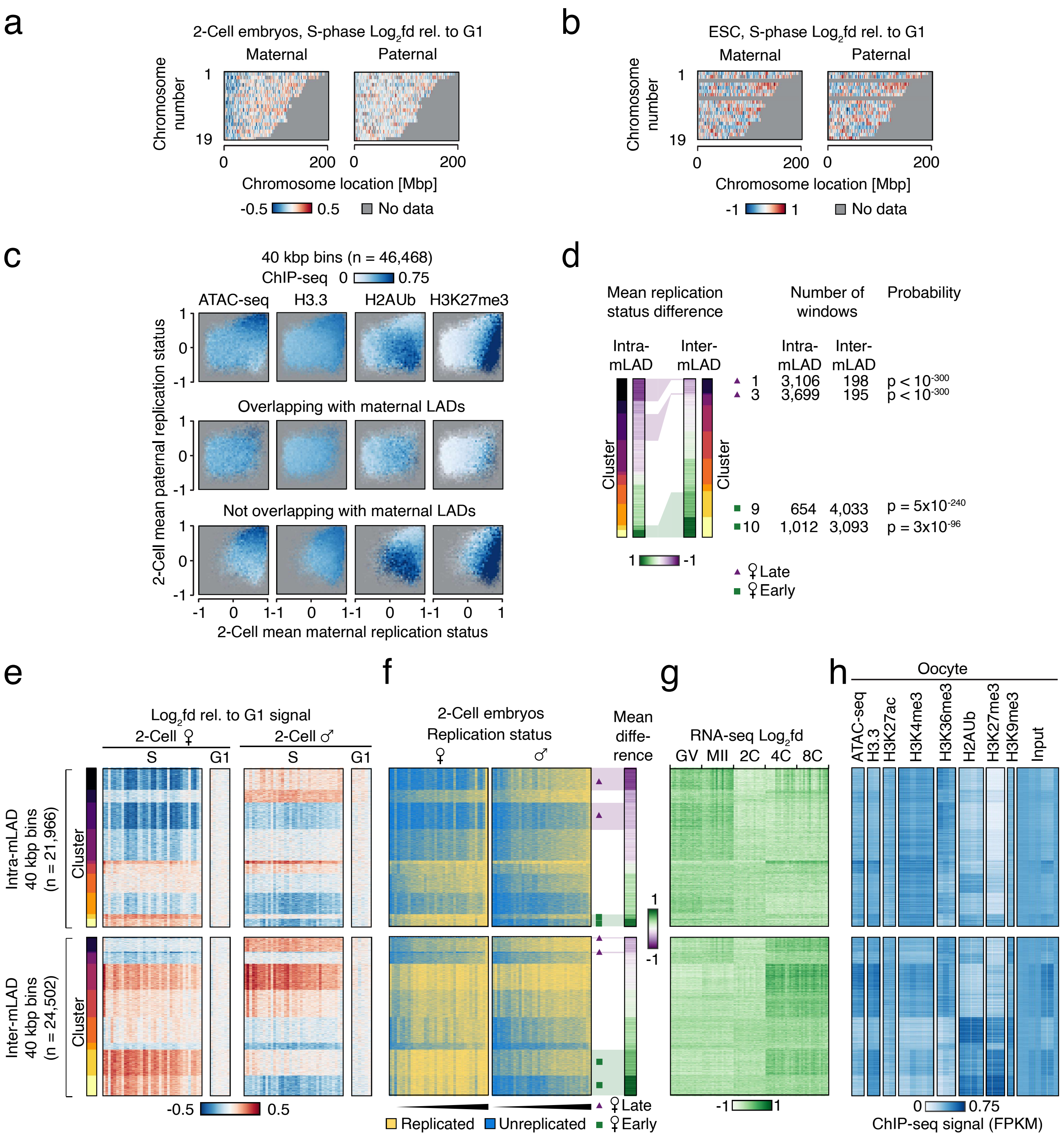


**Maternal early replication primarily occurred outside pericentromeres in non-overlapping LAD regions with high PRC marks.**

**a, b**) Heatmaps showing the normalized Repli-seq read densities in the maternal (left, n=27) and paternal (right, n=27) genomes of S-phase mouse 2-cell embryos (**a**) and ESCs (n=10) (**b**). X-axis position correspond to chromosome coordinates, and Y-axes correspond to autosome number. Aneuploidy was detected on chromosome 12 and 17 and therefore they were excluded.

**c**) Heatmaps showing the mean mouse oocyte histone marks, histone variant, and chromatin accessibility signal relative to mean maternal (X-axis) and paternal (Y-axis) binarized Replication Status. Values were quantified in 40 kbp windows in the entire genome (top), within (middle), or outside (bottom) LADs in the maternal genome of 2-cell mouse embryos.

**d**) Heatmaps showing the differences in the mean binarized Replication Status between the mouse 2-cell maternal and paternal genomes within (left) or outside (right) maternal 2-cell mouse embryo LADs. Clustering as in Figure **3c**. Green squares and purple triangles (as well as corresponding transparent overlays) indicate the extent of selected clusters constituting subpopulations of regions with a notably early or late replication of the maternal genome, respectively. Numbers indicate the counts of 40 kbp bins within each cluster inside and outside of LADs. P-values from $X^2$ tests Benjamini-Hochberg corrected for multiple testing. (*Continues on next page*)

**e-h**) Composite heatmaps showing a range of signal within (upper) or outside (lower) LADs in the maternal genome of 2-cell embryos. 40 kbp windows were k-means clustered according to the indicated S-phase signal from all individual maternal and paternal samples as in Figure **3c**, and clusters are ordered according to the mean difference between the maternal and paternal signal. From left to right: (**e**) Heatmaps showing the genome-wide signal enrichment and variation in individual Repli-seq samples from the maternal and paternal genome of S- and G1-phase 2-cell mouse embryos. (**f**) Heatmaps showing the genome-wide binarized Replication Status in individual Repli-seq samples from the maternal and paternal genome of S-phase 2-cell mouse embryos. Heatmaps showing differences in the mean binarized Replication Status between the maternal and paternal genomes. Green squares and purple triangles indicate selected clusters constituting subpopulations of regions with a notably early or late replication of the maternal genome, respectively. (**g**) Heatmaps showing transcript density difference from single mouse oocyte or embryo RNA-seq as well as (**h**) genome-wide histone mark, histone variant and chromatin accessibility signal from mouse oocytes. Signal was FPKM-normalized.
